# Digital Natives and Dual Task: Handling It But Not Immune Against Cognitive-Locomotor Interferences

**DOI:** 10.1101/723775

**Authors:** Dierick Frédéric, Buisseret Fabien, Renson Mathieu, Luta Adèle Mae

## Abstract

Digital natives developed in an electronic dual tasking world. This paper addresses two questions. Do digital natives respond differently under a cognitive load realized during a locomotor task in a dual-tasking paradigm and how does this address the concept of safety? We investigate the interplay between cognitive (talking and solving Raven’s matrices) and locomotor (walking on a treadmill) tasks in a sample of 17 graduate level participants. The costs of dual-tasking on gait were assessed by studying changes in stride interval time and its variability at long-range. A safety index was designed and computed from total relative change between the variability indices in the single walking and dual-task conditions. As expected, results indicate high Raven’s scores with gait changes found between the dual task conditions compared to the single walking task. Greater changes are observed in the talking condition compared to solving Raven’s matrices, resulting in high safety index values observed in 5 participants. We conclude that, although digital natives are efficient in performing the dual tasks when they are not emotional-based, modification of gait are observable. Due to the variation within participants and the observation of high safety index values in several of them, individuals that responded poorly to low cognitive loads should be encouraged to not perform dual task when executing a primate task of safety to themselves or others.

## Introduction

With the introduction of the early smart phone in the mid-1990s, the digital native generation was born into a world of internet and interactive screens [1, 2]. Interactive screen dual tasking, one could debate, was in actuality dual learning. Could such early development enhance the mental mechanisms creating a universal cognitive skill excluded to previous generations? Was this skill a new baseline at low cognitive loading and would standard gait changes occur as cognitive loading increased?

Technology based cognitive-motor tasks have become part of our daily lives. Driving with a maps application and walking while texting are standard examples. Yet, while some of us have witnessed the evaluation of the technology, for digital natives it is a baseline. Moppett defines digital natives as those born with the internet [1]. Additionally, while gait has been well studied, in terms of mental development could technology now to be influencing the process? We assume that dual task will impact gait variability, presumably leading to an increase of the latter. Fractal analysis, first applied to human walking by Haussdorff [3] is a powerful technique to assess the structural properties of gait variability. Hence, we resort to fractal analysis in the present work as well.

Previous studies have strived to understand the cognitive demands technology imposes as related to dual tasking [4]. Is reading the news on the smart phone different than reading a newspaper while walking? How talking on a phone vs. talking to a passenger is different in terms of driving [5]? These are all important research questions which should be used to inform the public of the safety trade-offs during dual tasking. In fact, these studies have aided new laws in several USA states to ensure driver vigilance [6]. What if a mind developed in a digital dual tasking world processes the demands differently than one adapted to it? To address this research question, the study would need to utilize well understood tasks and measures. Standards that are well established within both cognitive and motor-function domains.

The mental ability of dual tasking relies on executive function and working memory capacity both functions of the prefrontal cortex [7, 8]. When the two tasks interfere with one another, known as cognitive-motor interference, it is thought to be a proof of capacity limitation in cognitive abilities [9]. For safety considerations, this interference can become problematic if an individual’s cognitive resources are limited. In dual-task conditions, resources have to be shared between the two tasks and performances can decrease. Performance reductions from single- to dual-task conditions are often expressed as dual-task costs. The influence of expertise on cognitive-motor dual-tasking is predominant and experts are more successful than novices to keep up their performances [10]. As Dutke, Jaitner, Berse, and Barenberg indicate the results on dual tasking related to physical and cognitive demands is equivocal [8]. The selection and combination of the tasks greatly influence the outcome demands and resiliency [11]. Therefore, it is recommended that each task, especially the motor function, display a multifaceted output with high sensitivity to variation.

The objective of this study was to assess in digital natives, the impact of dual task-modified cerebral activity during talking and non-verbal reasoning on their spatio-temporal gait parameters. Special attention will be paid to variability of stride interval time and its organization at long-range to evaluate the cognitive-locomotor interferences.

## Material and Methods

### Participants and general protocol

#### Population

Participants were 17 masters students in Physical Therapy (9 ♂ and 8 ♀; age: mean±SD =21.5±1.6 years) from Haute Ecole Louvain en Hainaut (Montignies-sur-Sambre, Belgium). See Table 1 for individual features. This study was conducted in accordance with the declaration of Helsinki and all participants provided written informed consent under conditions approved by the Academic Ethical Committee Brussels Alliance for Research and Higher Education (acceptance nr: B200-2017-087). Average session duration was 45 minutes. Participants were not medicated and did not exhibit any neuromusculoskeletal, orthopaedic, respiratory, or cardiovascular disorders that could influence their gait. Participants could not have completed a Raven Progressive Matrices Test in the last six months. Participants were counterbalanced between the cognitive ability assessment and verbal task once the control condition was met.

**Table 1.** Individual characteristics of participants. A global description is given at the last line either under the form average± standard deviation or median [first quartile−third quartile].

|  | Age<br>(years) | Speed<br>(km/h) | Weight<br>(kg) | RAVEN |  |
| --- | --- | --- | --- | --- | --- |
|  |  |  |  | Score (/60) | Time (min) |
| 1 | 20 | 5.2 | 69.6 | 37 | 7 |
| 2 | 21 | 5.5 | 68.7 | 50 | 12 |
| 3 | 23 | 5.2 | 52.2 | 49 | 18 |
| 4 | 21 | 5.3 | 55.3 | 50 | 23 |
| 5 | 21 | 5.3 | 64.4 | 52 | 11 |
| 6 | 23 | 3.9 | 53.7 | 46 | 14 |
| 7 | 23 | 5.3 | 56.5 | 51 | 14 |
| 8 | 23 | 4.9 | 61.7 | 43 | 23 |
| 9 | 22 | 4.7 | 85.4 | 55 | 21 |
| 10 | 25 | 5.3 | 99.2 | 50 | 29 |
| 11 | 21 | 4.4 | 69.1 | 25 | 9 |
| 12 | 20 | 4.5 | 78.8 | 55 | 10 |
| 13 | 21 | 3.7 | 66.9 | 53 | 16 |
| 14 | 20 | 3.8 | 73.3 | 57 | 16 |
| 15 | 18 | 5.4 | 55.7 | 40 | 24 |
| 16 | 22 | 4.9 | 59.8 | 51 | 30 |
| 17 | 21 | 4.9 | 66.6 | 54 | 35 |
| | 21.5±1.6 | 4.8±0.6 | 66.9±11.9 | 50 [46 – 53] | 19±10 |

#### Physical Load

Participants walked on an instrumented treadmill (70 cm wide, 185 cm long) with an integrated force plate and an overhead safety frame (N-Mill, Motekforce Link, The Netherlands). They wore comfortable running shoes and a safety harness, and were asked to keep their eyes fixed straight ahead. Vertical ground reaction force (*F*_*v*_) and center of pressure (*CoP*) of each foot was recorded at a sampling rate of 500 Hz using the manufacturer’s software (CueFors 2, Motekforce Link, The Netherlands).

#### Control condition (CTRL)

Participants walked 10 minutes at their comfortable speed, determined during the familiarization procedure by the same experimenter (RM) and kept constant through the experimental duration. A minimal time of 10 minutes is needed to assess the fractal structure of walk, along the lines of [3].

#### Cognitive ability assessment (RAVEN)

Participants completed the modified Raven’s Progressive Matrices Test [13] while walking at their comfortable speed. The Raven’s test is a conventional measure of cognitive ability and was chosen to display participants fluid intelligence and a task related to pattern recognition during dual tasking [14]. It is made of 5 sets (A to E) of 12 items each. Participants walked the time necessary to complete the 60 items.

#### Verbal task (TALK)

Participants engaged in 10 minutes, non-emotional, conversation discussing their experience with ride sharing. This task intended to replicate a real time conversation with low cognitive workload during dual tasking.

### Data analysis

Stride interval (SI) time series were computed from heel strikes (during forward walking) or toe strikes (during backward walking) of the right foot identified on *F*_*v*_ − time histories and time series step width from maximal medio-lateral displacement of the *CoP* of two consecutive steps. At the completion of the session, three time series containing the values of the stride intervals in the different conditions were obtained for each subject. Typical plots are displayed in Fig. 1.

**Fig 1.**
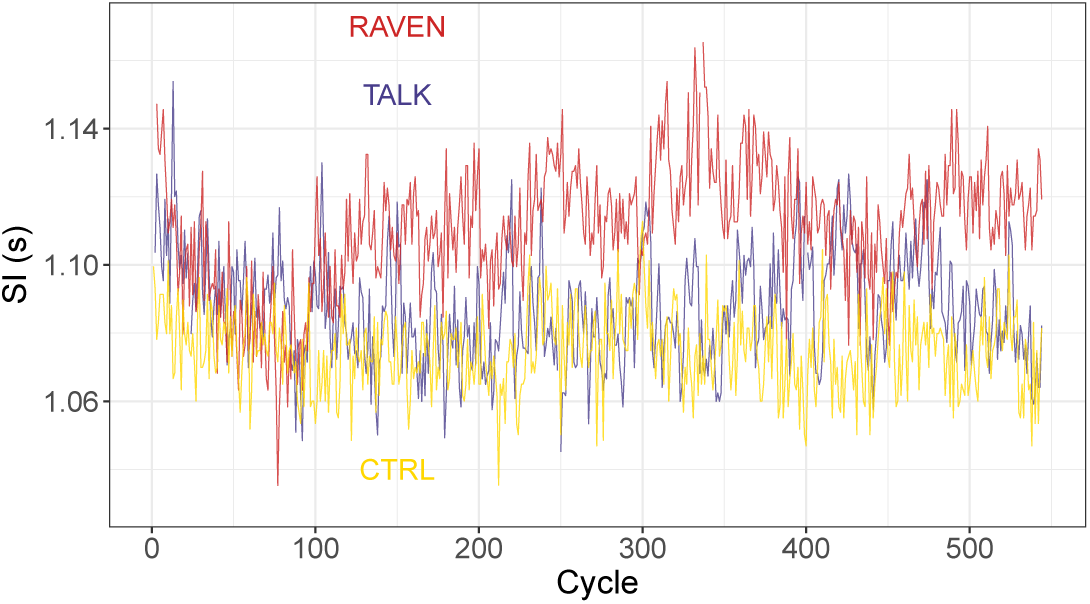
Typical plots of the SI time series obtained in the three conditions. The same subject (8) has been chosen.

Standard spatio-temporal indicators were also given by the instrumented treadmill’s software in each condition: The mean stride interval, denoted SI, step width, w, and dual stance time by cycle, DS. In the RAVEN condition, the score (number of correct items), the duration of a given item and the total time needed to complete the test were specifically recorded. The Belgium participant tables (1993) were utilized for grading the responses [13].

In addition to SI, w and DS, several variability parameters have been extracted from the recorded SI time series following the methods of Ref. [15]: The coefficient of variation, CV, the Minkowski fractal dimension, D, and the Hurst exponent, H, defined as follows. Let ***T*** be a SI time series containing the durations *T*_*i*_ of the successive cycles. The first variability indicator is *CV* = *SD*(***T***)*/SI*, estimating the relative amplitude of the SI fluctuations around the mean value. However, it provides no information on the temporal dynamics of these fluctuations. This is provided by two independent indices D and H [15]. The Minkowski fractal dimension D is computed by resorting to the box-counting method: if *N* (*ϵ*) is the number of square boxes of size *ϵ* needed to fully cover the plot of ***T*** (see Fig. 1), then *N* (*ϵ* → 0) *∝ ϵ*^−D^. D is related to the apparent roughness of the time series: The closer D is to 2, to more the SI relative fluctuations may be important from one cycle to another. Hence we relate D to the complexity of the time series dynamics. The Hurst exponent H is computed by using the Detrended Fluctuation Analysis (DFA) with a linear detrending and provides a diagnostic on the long-range trend of the time series [16]. The cumulated time series ***Z*** computed from ***T*** are divided into windows of length *l*. For each window, a local least squares linear fit is performed and the fluctuation function *F* (*l*), giving the average distance between the actual points and the fitted ones, is computed. The asymptotic relation *F* (*l* → *∞*) *∝ l*^H^ leads to H. A random process is close to *H* = 0.5, while long-range autocorrelated processes are such that 0.5 *< α ≤* 1 (*H >* 1 for unstable processes): An increase (resp.decrease) is likely to be followed by an other increase (resp. decrease) at long range. Time series with 0 *< H <* 0.5 are anticorrelated: An increase (resp.decrease) is likely to be followed by a decrease (resp. increase) at long range. H is regarded here as a predictability index of the time series: Strongly autocorrelated dynamics are such that the time series value at a given time is strongly dependent of its previous values, hence the time series is predictable. An innovative “safety index”, s, has finally been developed and computed by this lab. It is equal to a total relative change between the variability indices in the CTRL and in the perturbed (TALK and RAVEN) conditions:

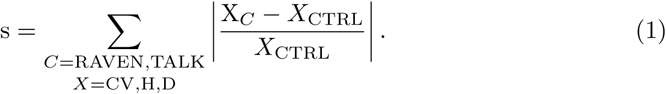

In the RAVEN condition, two extra parameters were computed: (1) The time shift Δ t is the difference between the duration of the first item of a set and the duration of the last item of the previous set. (2) The Pearson correlation coefficient, *r*, between the duration of a question and its number in a given set.

All the data analysis has been performed by R (v. 3.2.2) [17].

### Statistical analysis

Data were checked for normality (Shapiro-Wilk) and equal variance tests. A one-way repeated measures ANOVA with post hoc Holm-Sidak method was performed and used to examine the effects of condition (CTRL, RAVEN or TALK) on the computed parameters. The significance level was set at *p* = 0.05 for all analyses. In the case of a failed normality test, a one-way repeated ANOVA on rank with post hoc Tukey method has been performed (CV and DS are concerned). All statistical procedures were performed with SigmaPlot software version 11.0 (Systat Software, San Jose, CA).

## Results

### Differences between conditions

The condition has a significant influence on SI (*p* = 0.001), DS (*p* = 0.022), D (*p <* 0.001) and H (*p* = 0.011) as shown in Table 2. The post hoc analysis displayed in Table 3 reveals that the significant differences are driven by the differences between CTRL and the perturbed conditions TALK and RAVEN: (1) SI is significantly higher for the TALK (*p* = 0.02) and RAVEN (*p <* 0.001) conditions; (2) DS is significantly higher for the TALK condition (*p <* 0.05); (3) D is significantly lower for the TALK (*p* = 0.004) and RAVEN (*p <* 0.001) conditions; (4) H is significantly lower for the TALK condition (*p* = 0.003) than for the CTRL condition. No statistically significant difference is found between the TALK and RAVEN conditions for the parameters under study.

**Table 2.** Mean ± standard deviations values for the parameters computed in the three experimental conditions. Last line shows the *p* − value for the influence of the condition on the results.

| Condition | SI (s) | w (m) | DS (s) | CV | D | H |
| --- | --- | --- | --- | --- | --- | --- |
| CTRL | 1.103±0.069 | 0.098±0.017 | 0.293±0.029 | 0.0149±0.0094 | 1.696±0.105 | 0.891±0.166 |
| TALK | 1.119±0.076 | 0.097±0.023 | 0.297±0.030 | 0.0296±0.0307 | 1.558±0.230 | 0.753±0.153 |
| RAVEN | 1.121±0.079 | 0.093±0.020 | 0.296±0.030 | 0.0304±0.0334 | 1.500±0.138 | 0.812±0.216 |
| $p$ | **0.001** | 0.09 | **0.022** | 0.076 | **< 0.001** | **0.011** |

**Table 3.** Significance level (*p*–values) of the post hoc pairwise multiple comparisons between the different experimental conditions for the parameters analyzed. Statistically significant differences are written in bold font.

| $p$ | SI | w | DS | CV | D | H |
| --- | --- | --- | --- | --- | --- | --- |
| CTRL vs TALK | **0.02** | NS | **< 0.05** | NS | **0.004** | **0.003** |
| CTRL vs RAVEN | **< 0.001** | NS | NS | NS | **< 0.001** | 0.073 |
| TALK vs RAVEN | 0.687 | NS | NS | NS | 0.193 | 0.177 |

According to the experimenter in charge of the measurements (MR), all the participants blocked their upper limbs in some way (*e*.*g*. crossed the arms in the back) at some stage of the Raven’s test, and conversations were fluent in the TALK condition (*i*.*e*. walking on the treadmill did not prevent participants to have a normal conversation).

The above results are plotted in Figs. 2 and 3 as violin plots. It can be observed that the results’ dispersion is higher in the RAVEN and TALK conditions than in CTRL, even if the differences are not significative. The individual behavior of two participants has been added for completeness. Both participants have rather high values of *s* as it can be seen in Fig. 4, showing the safety indices computed from Eq. (1) for each participant.

**Fig 2.**
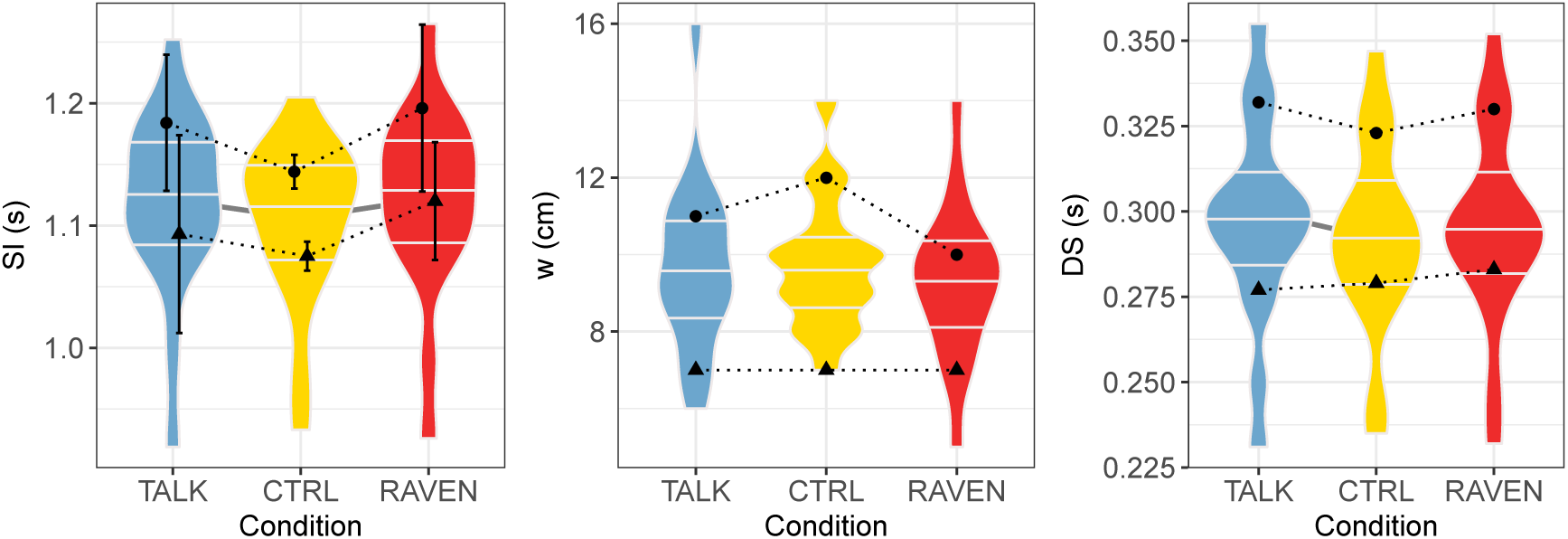
Violin plots of the spatio-temporal indicators SI, w and DS versus experimental conditions. The maximal and minimal data set the limits of the violin plots, and the median, first and third quartiles are also shown (horizontal bars). Significant differences between two conditions are marked by a gray line linking the concerned violin plots. The behaviour of two participants (3, circles, and 8, triangles) when going from CTRL condition to TALK or RAVEN conditions is displayed.

**Fig 3.**
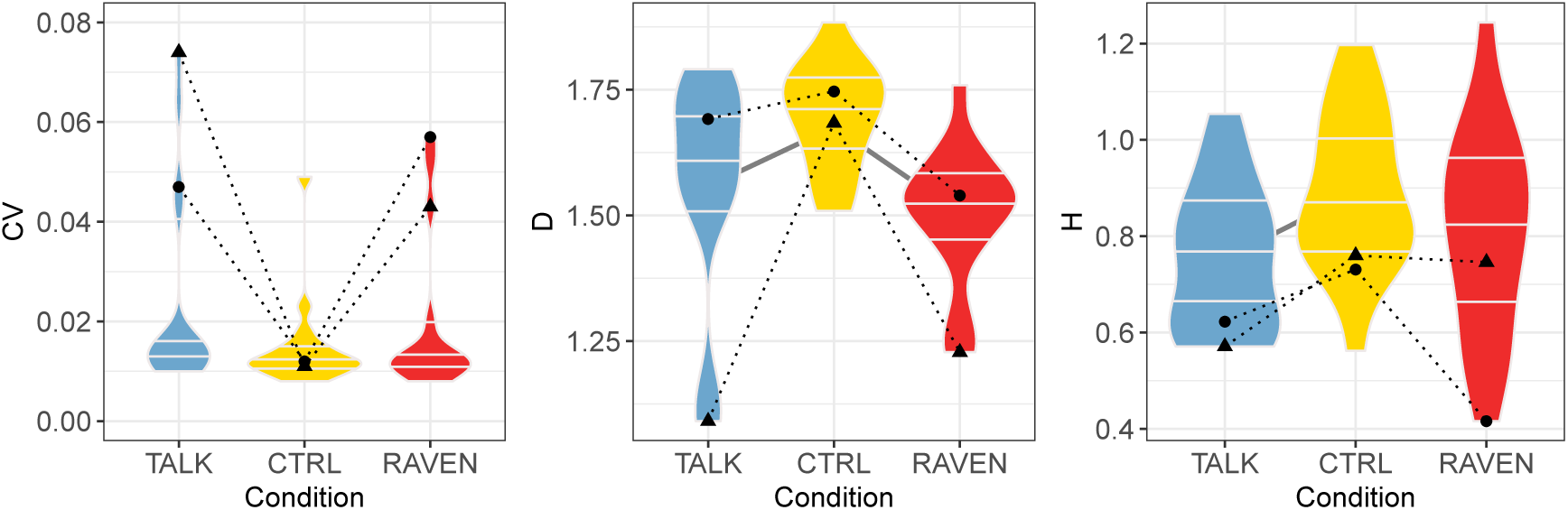
Same as Fig. 2 for the variability indicators CV, D and H.

**Fig 4.**
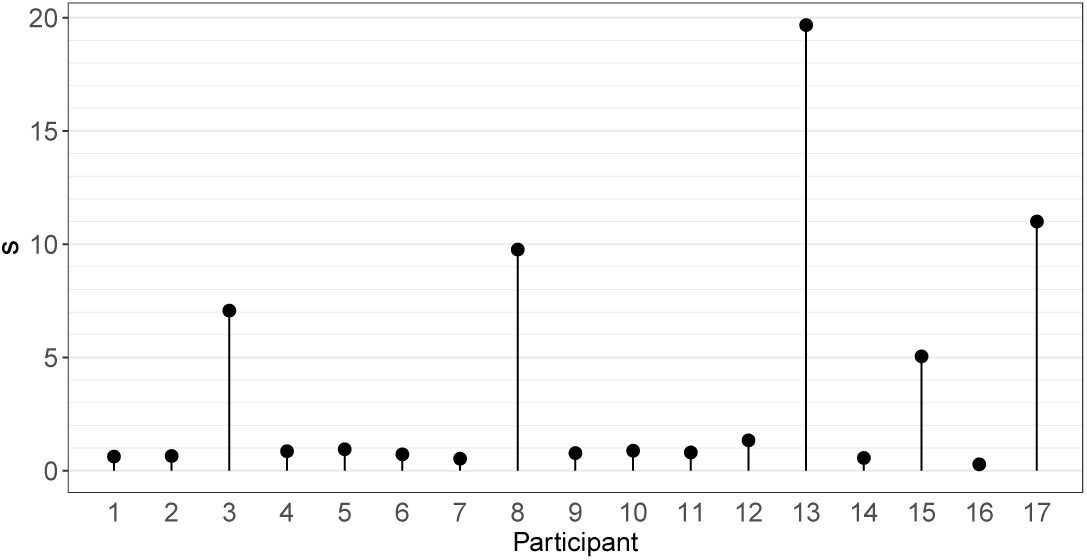
Safety index *s* computed for each participant.

### The RAVEN condition

Specific results are shown in Fig. 5 for the RAVEN condition. Several observations can be made: (1) For each set, more than 75% of the computed Pearson correlation coefficients between the duration and the question number are higher than 0.4. (2) More than 75% of the time shifts Δ*t* are negative between two consecutive sets. Hence we conclude that the first question of a set is solved more quickly than the last question of the previous set. From these observations it follows that the duration of questions vs the question number typically follows a sawtooth pattern, illustrated in Fig. 5.

**Fig 5.**
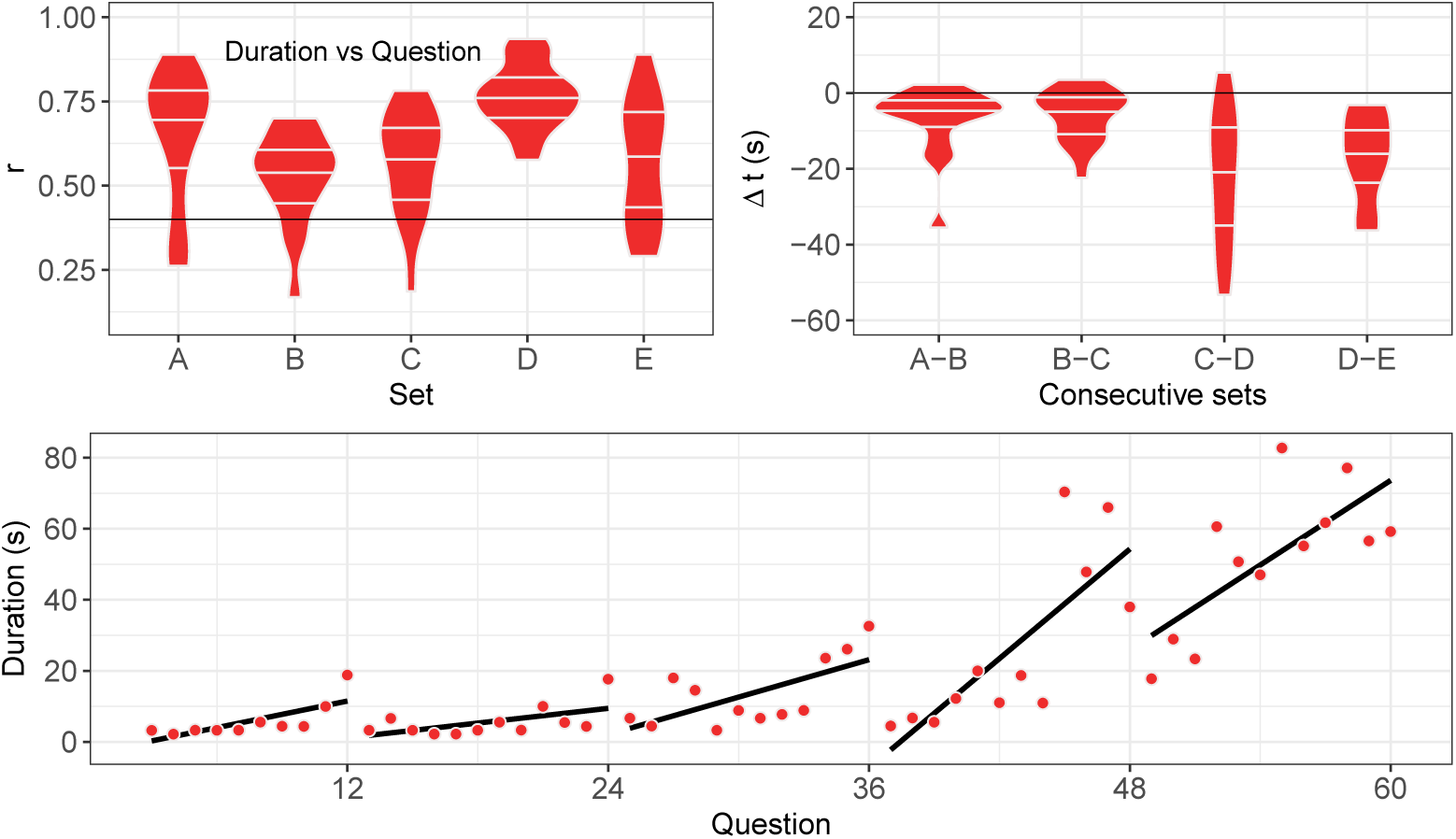
Upper left panel: Violin plots of the Pearson correlation coefficients *r* of the duration vs question number in a given set. The horizontal line *r* = 0.4 has been added to guide the eyes. Upper right panel: Violin plots of the duration differences Δ*t* between the last question of a given set and the first question of the next set. The horizontal line Δ*t* = 0 has been added to guide the eyes. Lower panel: Typical plot of the duration vs question number for the complete Raven test. Data of participant 8 have been plotted. Linear regressions of the duration vs question number within a given set are shown.

## Discussion

In previous studies [18–23], changes in different parameters such as walking speed, cadence, step length, double support time etc. were computed while performing secondary tasks to assess cost of dual-tasking on gait. Changes in gait parameters are believed to be associated with attentional capacity depending on the complexity of the secondary task [24]. Average activity in *α* and *β* frequencies was significantly modulated during both cognitive and motor interferences walking conditions in frontal and central brain regions, indicating an increased cognitive load during dual-task walking [25]. The coordination of multiple tasks is indeed an executive function, thought to be located in the prefrontal cortex [25]. The cerebellum regulates the cognitive and automatic processes of posture-gait control by acting on the cerebral cortex via the thalamocortical projection and on the brainstem, respectively [26]. The question raised in this study is the impact of dual task-modified cerebral activity on gait parameters, assessed by analysis of SI time series. We primarily computed variability indices, including the classical CV of SI time series and newer indexes exploring stride-to-stride temporal organization at long-range such as Hurst exponent (H) and Minkowski fractal dimension (D).

A first finding is that SI is significantly higher in both dual-task conditions: Participants are unable to decrease their speed as [19] treadmill speed is fixed, but they can increase their mean SI in order to slow down their pace. Such an increase has been observed by Bollens et al [21], although the difference was not statistically significant.

We also observed that a dual task tends to increase CV, *i*.*e*. pendular behavior of walk is controlled less efficiently. Both results globally fit into a cognitive-related motor interference model [27], where the dual task has a negative impact on the motor task. The dual task significantly decreases D in both conditions, which is coherent with the optimal variability framework [28]: Physiological signals – SI time series in the present work – show a maximal complexity in healthy participants, and any deviation from healthy state lowers it. As previously shown by our lab [15], D is a relevant measure of complexity in walking, hence we can state that SI time series complexity is lower because of the dual task. An intuitive argument, related to capacity-sharing model, is that participant’s mind is “busy” while talking or solving Raven’s matrices, resulting in a lower ability to handle afferent signals needed to optimally adapt their walk and eventually in a lower D. We can safely assume that adding a cognitive load during gait in digital natives has noticeable effects on SI and its stride-to-stride temporal organization. However, the consequences are dependent on the nature of the cognitive load since modifications were not similar in TALK and RAVEN conditions.

Walking while talking can be seen as an everyday example in which walking is combined with a cognitive task. Such paradigms involve executive functioning [29–31]. Both executive function and gait performance have been shown to be closely associated with frontal lobe function, whereas walking is specifically connected with activity in the primary motor cortex, the supplementary motor area, and also the premotor cortex [32–34]. This increased activity particularly in prefrontal areas appears important to resolve both tasks properly. In digital natives, TALK condition has a greater cost of dual-tasking than RAVEN: DS and H are significantly modified in TALK with respect to CTRL condition, but no equivalent effect is observed in the RAVEN condition. An increase in DS has been previously found by Ebersbach et al. [35] during the simultaneous realization of a fine motor and a memory task. The decrease of H suggests that talking induces random perturbation of walking mechanism, leading to a less predictable SI time series at long-range.

To our knowledge, it is the first time that Raven’s Progressive Matrices Test were used as a cognitive load during walking. This task assesses fluid intelligence, characterized by the capacity of quickly abstracting rules in a versatile manner. A previous study showed that the locomotor system is adaptive enough to complete a working memory task (OSPAN) while not compromising gait performance when walking at a self-selected pace [36]. Fluid intelligence has been thought to rely, at least partly, on working memory function [37, 38]. However, Engle et al. (1999) has found that fluid intelligence is only weakly (*r <* 0.4) correlated with OSPAN in young adults [39]. Therefore, the influence of fluid intelligence when walking has never been fully explored up to now. Raven’s matrices were used both as a cognitive load and a tool for assessing cognitive performance during the dual task. The foremost pronounced changes in cerebral activation patterns has been showed to occur during the second set of the test [40] and concerned associative somatosensory area and Wernicke’s area that is known to play a crucial role in cognitive processes related with synthesis and analysis of information. Interestingly, set B shows the poorest correlation between duration and question number. According to a Rasch model study [41], sets B and E are not unidimensional. Set E coherently shows the maximal dispersion of the results concerning the duration vs question correlation coefficient: Individual differences are maximal in that set. We noted that participants stopped moving their arms in RAVEN condition: It shows an effort to provide more resources to the task.

The definition of the safety index has been chosen along the lines of the optimal variability framework: The more the variability indices CV, D and H are close to the value in CTRL condition the more s is small. Our findings show that 5 on 17 participants (nr 3, 8, 13, 15 and 17, see Fig. 4) have values well larger than the others, suggesting that the dual-task of walking while talking or completing Raven’s Progressive Matrices Test impacts executive function and influences gait to such a degree that it may compromise safety. Altered stride-to-stride variability is indeed related to risk of fall, as reviewed in [42]. It is clear that not enough participants took part to the study to identify the factors leading to high s but even at this stage it appears that digital natives are not as immune against dual task as expected, as already stressed in the domain of education [43].

### Future work

While we are confident of the resolution within the dataset, increasing range selection of fluid intelligence would aid in determining if the variability across participants reflects an accurate range in safety index. Additionally, including a higher tasking verbal test to reflect the range found within the Raven’s test would be beneficial. It was important at this initial stage to not impede natural walking motion but given this baseline future data collections could replicate external studies by allowing for participants to hold smartphones while performing the tasks. We hope to present such a study in the future.

